# Exploring Spike-Dependent and ACE2-Independent Viral Entry into Salivary Epithelial Cells in the Absence of ACE2

**DOI:** 10.1101/2025.05.19.654917

**Authors:** Caitlynn Barrows, Simon Young, Mary Farach-Carson

**Affiliations:** UTHealth Houston School of Dentistry, Katz Department of Oral and Maxillofacial Surgery; UTHealth Houston School of Dentistry, Department of Diagnostic and Biomedical Sciences

**Keywords:** Salivary gland, ACE2, SARS-CoV-2, spike protein, COVID-19

## Abstract

Salivary gland infection by SARS-CoV-2 requires viral entry via routes and mechanisms that remain unresolved. This study examined the expression of the angiotensin-converting enzyme 2 (ACE2) receptor in salivary tissues and basal cell-derived human salivary progenitor cells (hS/PCs), an unstudied potential entry point for SARS-CoV-2. Multiple detection modalities, including immunocytochemistry, western blotting, flow cytometry and RT-PCR, demonstrated a consistent lack of ACE2 protein and transcript in both tissue specimens and primary salivary epithelial cells. Antigen retrieval at pH 9 was determined to be optimal for immunodetection protocols, yet ACE2 remained undetectable. Small intestine tissue served as a positive control, confirming the validity of the methods and reagents we used. Considering there can be other receptors for SARS-CoV-2, flow cytometric analyses demonstrated that recombinant SARS-CoV-2 spike protein failed to bind to salivary epithelial cells, in contrast to HEK293 cells engineered to overexpress ACE2, which showed robust spike binding. These findings strongly support our conclusion that salivary cells do not serve as major targets for SARS-CoV-2 infection via ACE2 or spike protein, whether through direct exposure to viral particles in ductal fluids or via access to basal cells across the basement membrane.

## 1. Introduction

In early 2019, Severe Acute Respiratory Syndrome Coronavirus 2 (SARS-CoV-2) broke out in Wuhan, China [1] leading to the global pandemic known as COVID-19. During its spread from Wuhan across the globe, SARS-CoV-2 acquired significant mutations associated with increased virulence and evasion of the immune system [2]. The most significant of these mutations occurred in the viral spike protein, the site of contact and entry of the virus into human cells [3]. The transmission mode was identified as person-to-person through contact with aerosols and droplets containing the respiratory virus [4]. This transmission method indicates that the upper respiratory system specifically has high involvement in viral spread. Given the similarities between SARS-CoV and SARS-CoV-2, the Angiotensin Converting Enzyme 2 (ACE2) receptor was quickly identified as the primary site for SARS-CoV-2 binding and became the accepted receptor for viral entry [5], [6]. Consequently, several studies explored the tissue locations of ACE2 to identify the potential impact of SARS-CoV-2 on various organs and additional possible sites of direct infectivity beyond the lung [7], [8]. One study implicated the salivary glands as a potential site of infectivity based on bulk RNA sequencing data obtained from publicly available databases including GTEx and Fantom 5, which indicated that the salivary glands had similar transcript levels of ACE2 as compared to the lung [9]. At the time of publication, the nasopharyngeal swab was the gold standard for SARS-CoV-2 detection and remains the preferred way of testing, although saliva-based diagnostics have been developed[10], [11].

The presence of active viral particles in saliva [12] suggested a role for salivary glands in disease transmission through expelled salivary droplets, in addition to those originating in the nasopharynx or respiratory system. If direct infection of salivary glands occurred, then the pathway by which the virus reaches the salivary epithelium, where any receptors associated with infection would be located, must be deduced. Given their locations and abundance throughout the oral cavity, the 800–1000 minor salivary glands throughout the oral cavity were given initial consideration as sites of primary infection, and some reports demonstrated presence of viral particles present in these minor glands [13]. Despite this observation, it is not entirely clear how the SARS-CoV-2 virus, if present in the oral cavity, would reach the surface and infect these minor glands in titers high enough for general infection. Minor glands present beneath the oral mucosa (10% of total salivary production), lack long, named ducts like those in the major glands, and only possess very short excretory ducts 10-50 μ in diameter that produce a mucous-rich secretion that protects the glands from general infection[14], [15]. These microscopic ducts open directly onto the mucosal surface, often within or just beneath the epithelium where they lubricate the oral cavity. The SARS-CoV-2 virus, 100-140 nm in diameter covered with 20 nm spike proteins [16], is small enough to penetrate the excretory ducts, but would need to actively move through mucous secretions in a direction retrograde to fluid flow from the minor glands.

Humans possess three pairs of major salivary glands, parotid, submandibular and sublingual (90% of saliva production), connected to the oral cavity through major ducts [17], [18]. The largest gland, the parotid, is connected to the oral cavity through Stensen’s duct, typically 5–6 cm in length and 1-3 mm in diameter. The gland opens opposite the second maxillary molar, via a papilla in the buccal mucosa, and opens into the oral cavity. The parotid gland is serous, producing copious amounts of watery, amylase-rich saliva. The submandibular gland is connected to the oral cavity through Wharton’s duct, approximately 5 cm in length and 1-3 mm in diameter, and produces a mixed saliva that is the product of both serous and mucous acinar cells. Wharton’s duct opens at the sublingual caruncle, under the tongue. The sublingual gland is the most mucous-rich of the major glands, and does not rely on a single large duct like the parotid and submandibular glands. Rather, 8-20 small, shorter ducts of Rivinus (1-2 mm) open directly onto the floor of the mouth alongside the sublingual fold. Bartholin’s duct (1-2 cm in length), when present, may join Wharton’s duct or have a nearby separate opening. Unstimulated saliva in normal salivary glands flows at rates of ∼0.3–0.5 mL/min, whereas stimulated flow reaches rates of ∼1.5–2.0 mL/min, with daily production reaching up to 1.5 liters per day [19], [20]. SARS-CoV-2 virus particles present in the oral cavity would need to move against a significant flow of fluid, then penetrate the mucous-lined walls of the ducts [21], to enter and infect the secretory acinar component of the major salivary glands.

The structure of the major exocrine salivary glands includes terminal end buds consisting of serous and mucosal acini, present in various proportions in different glands, that produce the saliva. Saliva produced by acinar cells is transported and buffered by an organized network of ductal cells before being secreted into the oral cavity. As the larger ducts patent in the oral cavity approach the terminal endbuds where saliva is produced, they narrow and branch. Larger interlobular ducts give way to small intralobular/striated ducts and finally to intercalated ducts that conduct saliva from secretory acini to striated ducts. The intercalated ducts of the major salivary glands are the smallest ducts in the ductal system and serve as the initial conduit for saliva produced by acinar cells. Intercalated ducts play important roles in both ion transport and antimicrobial defense [18], [22]. Their cellular architecture, including the presence of basal cells, consists of a simple, contiguous cuboidal epithelium with regularly spaced basal cells that rest beneath the cuboidal cells and abut the basement membrane separating epithelium from the underlying stromal compartment. The basal cell population is key for ductal regeneration and repair, and expresses markers including K5, K14, p63 [23]. Our laboratories have studied this cell population for years, having extracted them from both major and minor glands [24], [25] and shown them to be a rich stem, progenitor population for tissue engineering and regeneration that we term human stem/progenitor cells (hS/PCs) [23], [26]. Relative to studies of SARS-CoV-2 infection of salivary glands, these cells are of high interest as their infection could have the ability to be propagated throughout the gland, unlike that of the terminally differentiated cuboidal ductal cells lining the ducts. Reports have indicated that basal cells present in the ducts express both ACE2 and TMPRSS2, the co-receptor for SARS-CoV-2 [27]. Because of their proximity to stroma and the vascular system of the salivary gland, salivary basal cells could conceivably be infected by viral particles in circulation spreading from other sites of major SARS-CoV-2 infection. Entry through this route would be unlikely to involve ACE2, but could involve an alternate receptor, integrins α5β3 and α5β1 [28], [29], demonstrated to facilitate entry of SARS-CoV-2 [30], presumably from circulation through a compromised basement membrane[31]. This integrin has previously been shown to be expressed in salivary epithelial cells [32], [33].

In this work, we set out to determine systematically where ACE2 is localized in the complex structures of the salivary gland, which cells could be targeted by SARS-CoV-2 via an ACE2-dependent pathway and whether basal cells could be recognized by spike protein. This information could shed new light on the routes to infectivity of salivary cells by coronaviruses.

## 2. Materials and Methods

### 2.1 Cell Culture

Primary cells were isolated from salivary glands. All procedures for tissue collection followed approved Institutional Research Board guidelines at UTHealth Houston. Patients provided informed consent, and all sample collections were deidentified except for sex and age. Human parotid and human minor salivary glands were processed as previously described[24], [25]. Briefly, salivary glands were washed in 1X PBS without calcium and magnesium. Glands then were washed in cold DMEM/F12 (Gibco) supplemented with 1% v/v penicillin-streptomycin, 1% v/v amphotericin B, 1% v/v betadine solution (from 10% povidone iodine stock). A final wash used cold DMEM/F12 supplemented with 1%v/v penicillin-streptomycin, 1% v/v amphotericin B. All solutions were sterile filtered through a 0.22 μm PES filter. Glands were minced with scissors to create a slurry and plated in a T-75 flask with 5 ml of William’s Medium E without L-glutamine (Sigma, W4128) supplemented with 1 mg/mL albumin from human serum (Sigma, A1887), 1% v/v CTS™ GlutaMAX™-I Supplement (Gibco, A12860-01), 1% v/v penicillin-streptomycin, 1% v/v insulin-transferrin-selenium (InVitria, 777ITS032), 0.1 μM dexamethasone, and 10 ng/mL human epidermal growth factor (Gibco, PHG0311). Cultures were left for 5 days to allow for full adherence of explants to the cell culture flask before media was changed. After 5 – 7 days, the first passages were conducted with 0.125% (v/v from stock) trypsin-EDTA.

Additional cell lines. HEK293T (ATCC, CRL-3216) and ACE2-HEK293T (Genecopoeia, SL221) were cultured in DMEM high glucose (Corning, 10-013-CV) supplemented with 10% v/v FBS and 1% v/v penicillin-streptomycin. ACE2-HEK293T medium was supplemented with 100 μg/mL hygromycin-B. CACO2 (ATCC, HTB-37) cells were cultured in EMEM (ATCC, 30-2003) supplemented with 20% v/v FBS and 1% v/v penicillin-streptomycin. All cultured cells were maintained in an incubator at 37ºC with 5% CO2.

### 2.2 Tissue Processing and Immunohistochemistry

Small intestine sample and salivary parotid and minor gland samples were placed in 10% v/v formalin for 24 hours, paraffin embedded and sectioned (5 μm sections) by the Oral and Maxillofacial Pathology Core. Deparaffinization was conducted with Histo-Clear® (Electron Microscopy Sciences, 64110) followed by rehydration in decreasing concentrations of ethanol. Three conditions were tested, no antigen retrieval, antigen retrieval with AR6 Buffer (Akoya Biosciences), or antigen retrieval with AR9 Buffer (Akoya Biosciences). Heat-mediated antigen retrieval steps were conducted in a Biogenex microwave at 95ºC for 15 minutes. Slides were cooled to room temperature and washed with 1X PBS. Sections were permeabilized with 0.3% Triton x-100 v/v for 30 minutes followed by a one hour blocking step with 10% normal goat serum and 0.3% triton x-100. Subsequently, sections were incubated in primary antibodies overnight at 4ºC: ACE2 monoclonal antibody (ProteinTech, 66699, 1:100), ACE2 monoclonal antibody (Abcam, 272500, 1:100), ACE2 polyclonal antibody (Abcam, 15348, 1:1000), Pan-cytokeratin (Abcam, AB234297, 1:100) or α-SMA (Abcam, AB7817, 1:100). Following the primary incubation, sections were washed three times with PBS and incubated with secondary antibodies: goat anti-rabbit 488 (Invitrogen, A11008, 1:1000), goat anti-mouse 568 (Invitrogen, A11004, 1:1000), goat anti-rabbit 568 (Invitrogen, A11036, 1:1000), or goat anti-mouse 488 (Invitrogen, A11029, 1:1000). DAPI was used to visualize the nuclei (Thermo Scientific, 62247, 1:500 of 1mg/mL stock). Confocal fluorescence microscopy imaging was performed on the Nikon A1R/MP.

### 2.3 RNA Isolation and qPCR

Cells were lysed with 1ml of TRIzol® LS Reagent (Life Technologies, 10296028) and collected in Eppendorf tubes. After lysis, the Direct-zol RNA mini prep kit was used (Zymo Research, R2052) according to the manufacturer’s protocol. Briefly, equal volumes of 100% ethanol were used and each mixture was transferred to the spin column. Samples were spun and treated with DNAse 1. Washed samples were eluted with nuclease free water. RNA purity and concentration were measured using a nanodrop (Thermo Scientific Nanodrop™ 2000). Following RNA isolation, a two-step PCR protocol was conducted. First, cDNA was generated using qScript cDNA SuperMix (Quanta Biosciences, 95048-1000) according to the manufacturer’s protocol. Briefly, RNA (1 μg) was added to 4 μl of qScript cDNA; water was added to balance 20 μl. The cDNA was made in a thermocycler with the following parameters: incubate at 25ºC for 5 minutes, 42ºC for 60 minutes, then 85ºC for 5 minutes. After cDNA was constructed, the PCR reaction was conducted with the PerfeCTa SYBR® Green SuperMix (Quanta bio, 95053) with human ACE2 primers [34] forward 5’ TCCATTGGTCTTCTGTCACCCG 3’ and reverse 5’ AGACCATCCACCTCCACTTCTC 3’ and GAPDH primers 5’ TTGAGGTCAATGAAGGGGTC 3’ and reverse 5’ GAAGGTGAAGGTCGGAGTCA 3’.

### 2.4 Protein Isolation and Western Blot

Cells were lysed with RIPA Buffer that included EDTA and protease and phosphatase inhibitors. A BCA assay was conducted and read on a Biotek Cytation 5 to determine protein concentrations and total protein (20 μg) was added. When using protein isolated from the overexpressing ACE2 HEK293T cell line, only 5 μg of total protein was added. Protein was added to Laemmli buffer and boiled at 100º C for 5 minutes. Samples were electrophoresed on 10% Bis-Tris gels for 1 – 1.5 hours with MOPS running buffer. Following the gel separation, a wet-transfer was carried out to transfer protein to a nitrocellulose membrane. The transfer was performed with 1X TG buffer at 50 V for 5 hours with an ice block at 4ºC. Nitrocellulose membranes were washed 3X with 1X TBST buffer and blocked for one hour at room temperature with 3% BSA (Fraction V, heavy metal reduced). Primary antibodies were incubated overnight at 4º C: ACE2 monoclonal antibody (Abcam AB272500 1:1000) and GAPDH (Invitrogen, PA1-987, 1:10,000). After 3 washes, secondary goat anti-rabbit HRP (1:50,000) was administered for one hour. Finally, an additional 3 washes were completed and then HRP chemiluminescent substrate was used and the blot was exposed to x-ray film, developed on a Konica Minolta SRX-101A, and then scanned on an Epson Perfection V30.

For whole tissue extracts, tissue was homogenized with a dounce homogenizer on ice with RIPA buffer, EDTA, and protease/phosphatase inhibitors. Samples were spun at 10,000 g for 15 minutes and the clear supernatant layer was extracted. Samples were aliquoted and stored at -20o C until all collections were completed. The BSA assay for protein concentration and remaining western blot protocol followed the cellular protocol. The small intestine fresh frozen positive control sample was purchased from BioIVT. The whole tissue blots were imaged on a ChemiDoc™ MP imaging system.

### 2.5 Flow Cytometry

ACE2: Cells (salivary primary cells or ACE2-HEK293T) cells were trypsinized and resuspended in PBS with calcium and magnesium. Cells were prepared and then blocked with 10% goat serum in 1X PBS followed by incubation with the ACE2 monoclonal antibody (Ab272500, 1:500). Cells were washed three times with 1X PBS and the secondary antibody goat anti-rabbit 488 was used.

Spike Binding: Live/Dead Fixable Aqua was used to identify live cells (Invitrogen, L34965). Cells were treated with either Alpha spike protein (R&D Systems, AFG10796) or Omicron BA.2 (R&D Systems, AFG11109) for 30 minutes followed by 3 washes with 1X PBS. Samples were kept on ice for the duration of the staining. Data was acquired on a BD Biosciences LSRFortessa™ and analyzed with FlowJo software v10. Cells again were washed 3X and analyzed as described above.

## 3. Results

### 3.1 Tissue Level Examination of ACE2 in Minor Salivary Glands and Parotid Tissue

In the first set of studies using fluorescence-based immunocytochemistry on fixed tissue, we tested a series of three antibodies that have been used in various laboratories for detection of ACE2 protein in fixed tissues from both major (parotid) and minor salivary gland tissues. Small intestinal tissue served as the positive control. Because previous studies have inconsistently reported a presence or absence of ACE2 in salivary tissues, we systematically compared the signal obtained without antigen retrieval and after antigen retrieval at pH 6 and pH 9. In each case, tissues were counterstained with either α-smooth muscle actin (α-SMA) or a pan-cytokeratin (pan CK). In salivary tissues, α-SMA serves as an excellent marker for myoepithelial cells surrounding each salivary acinus, while pan CK should stain all epithelial cells.

Figure 1 shows the results of the immunocytochemical analysis. Without antigen retrieval (Figure 1A panels, bottom row), no ACE2 labeling was observed in the small intestine positive control. This was the case for all three ACE2 antibodies, one rabbit polyclonal (PAb15348) and two mouse monoclonals (MAb272500, MAb66699) (Figure 1A). Likewise, without retrieval, no ACE2 was detected in parotid gland nor two minor salivary gland specimens (Figure 1A, top three rows). In the small intestine not subjected to antigen retrieval, α-SMA showed scattered and discontinuous staining in the lamina propria (red color) with two of the antibodies used, and pan-CK was not observed. Modest, diffuse staining for α-SMA was seen in the acinar, but not ductal, regions of both parotid and minor salivary glands. Given these negative findings, all subsequent studies used antigen retrieval.

**Figure 1.**
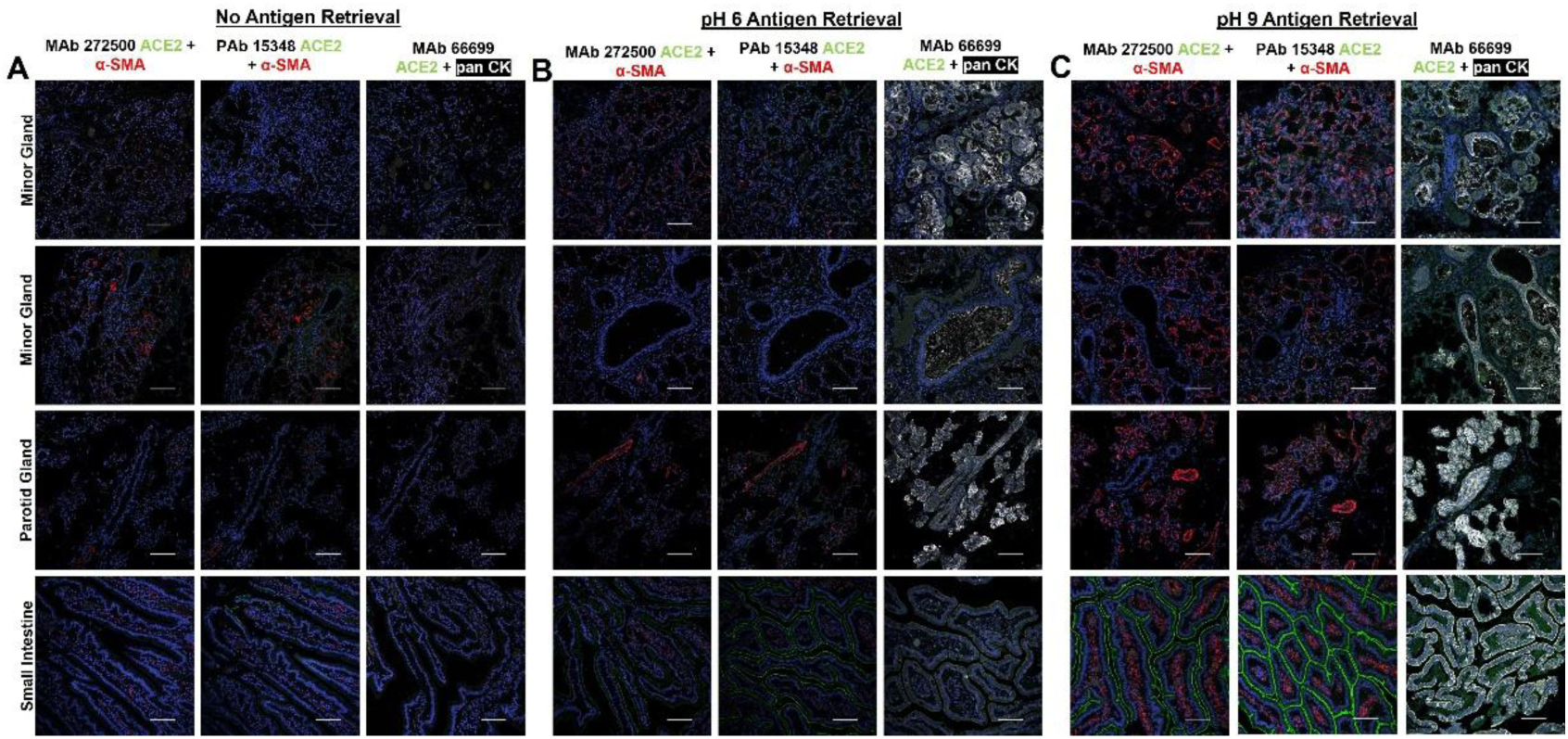
Immunofluorescence labeling in salivary glands and small intestine tissue sections shows lack of ACE2 in major and minor salivary glands. Three anti-ACE2 antibodies (and three conditions were used for detection of ACE2 with small intestine serving as the positive control. (A) ACE2 detection without antigen retrieval. (B) ACE2 detection after pH 6 antigen retrieval (C) ACE2 detection after pH 9 antigen retrieval. In all panels ACE2 is labeled green and α-SMA red or pan-CK white. DAPI is used to visualize nuclei in blue.

In the studies shown in Figure 1B, all fixed, serial sections underwent antigen retrieval with a pH 6 based antigen retrieval, AR6 buffer. After retrieval, the polyclonal antibody (Mab15348) treated specimens showed robust signal on the apical side of the small intestine epithelium (Figure 1B, bottom row). Similarly, the monoclonal antibody (ab272500) treated small intestine sections showed fainter, but still positive, signal in the same location. Sections treated with the second monoclonal antibody (MAb66699) showed little or no positive signal in the small intestine even after pH 6 antigen retrieval. After retrieval at pH 6, α-SMA was visible in the lamina propria of the small intestine and pan-CK (white) had a slightly positive signal in the epithelial compartment. For the salivary gland sections examined (Figure 1B, top three rows), no staining for ACE2 was seen with any of the three anti-ACE2 antibodies. Signal for α-SMA was observed in both acinar and vascular cells with robust staining of both acinar and ductal epithelial cells seen with pan-CK.

To determine if a pH 9 based antigen retrieval would yield clearer results, a pH 9, AR9 buffer was used. After this treatment, both MAb272500 and PAb15348 antibodies for ACE2 showed strong and robust signal (green) on the apical surface of the small intestinal epithelium (Figure 1C, bottom row). Detection for α-SMA showed strong signal localized to the lamina propria compartments of the small intestine not overlapping the staining for ACE2. Anti-CK staining of the epithelial compartment also was strong and uniform after retrieval at pH 9. Examination of salivary gland sections after retrieval at pH 9 (Figure 1C, upper three rows) showed strong staining for α-SMA in the myoepithelial component of both salivary parotid and minor glands, as well as vascular elements. No staining of salivary ducts was observed. Pan-CK staining showed strong expression in the epithelial compartment including both acinar and ductal cells present in parotid and minor salivary glands. In contrast, even under conditions where all three anti-ACE2 antibodies produced strong signal in the positive control, no ACE2 staining was visible in either the parotid or minor salivary glands. Signal was absent in both acinar and ductal compartments, even when these distinct structures were clearly visible in the specimens, providing strong evidence that there is likely to be very little ACE2 present on the surfaces of exocrine salivary cells.

To verify if this lack of ACE2 protein in salivary glands was true using an alternative method, whole tissue lysates were analyzed by western blot. As shown in Figure 2, no positive signal for ACE2 was seen in protein extracted from two parotid glands and three minor glands. In contrast, positive signal was seen for both small intestine lysate and a cell line, ACE2 HEK293T, constitutively producing ACE2 from a constitutively expressed transgene.

**Figure 2.**
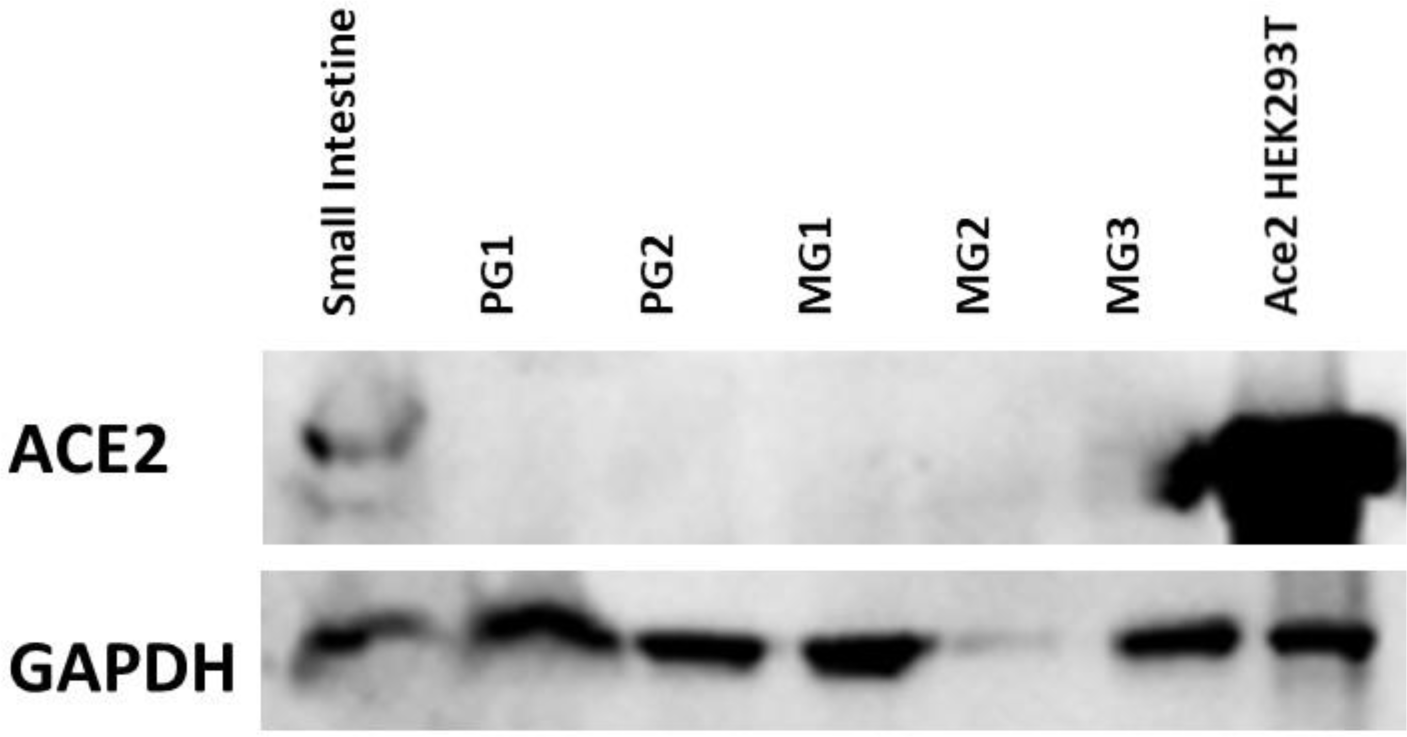
ACE2 detection in protein extracts from small intestine, major and minor salivary glands and ACE2 HEK293T cells. Western blotting for ACE2 (upper bands) and GAPDH (lower bands) in small intestine (left), and five salivary glands from different patient donors, including two parotid and three minor salivary glands. ACE2 HEK293T cells are shown on the right and were used as an additional positive control along with small intestine. No signal was seen in any salivary tissue extract.

### 3.2 Cellular Examination of ACE2 Transcripts and Protein in Salivary Basal Cell-Dervived hS/PCs

Given our inability to identify ACE2 protein in salivary tissue by immunocytology under the best conditions, we next sought to determine using different techniques if ACE2 transcripts or protein were present in expanded cultures of pluripotent, primary salivary epithelial cells. Salivary cells derived from minor and major glands of multiple patients were examined using qPCR and validated primers specific for human ACE2 transcripts. ACE2-HEK293T cells served as the positive control and expressed high levels of transcript, ∼500 fold increase over CACO2 cells (Figure 3A). CACO2 cells also expressed ACE2 transcripts, but at significantly lower levels than ACE2 HEK293T’s (Fig 3A). Four salivary primary cell populations from different sexes andbackgrounds were tested independently (M88C, F71AA, F69b, M32). None of these cultured lines from major or minor (M32) glands were found to express detectable levels of ACE2 transcripts (Figure 3A). Considering that expression of ACE2 by salivary cells might be a result of chronic inflammation during SARS-CoV-2 infection, we next set out to determine if we could force ACE2 expression through exposure to the inflammatory cytokines TNFα, IFNγ, or a combination of the two. After 24 hours and 72 hours of treatment, none of the treatments induced expression of ACE2 transcripts (Figure 1 Supplemental Figures).

**Figure 3.**
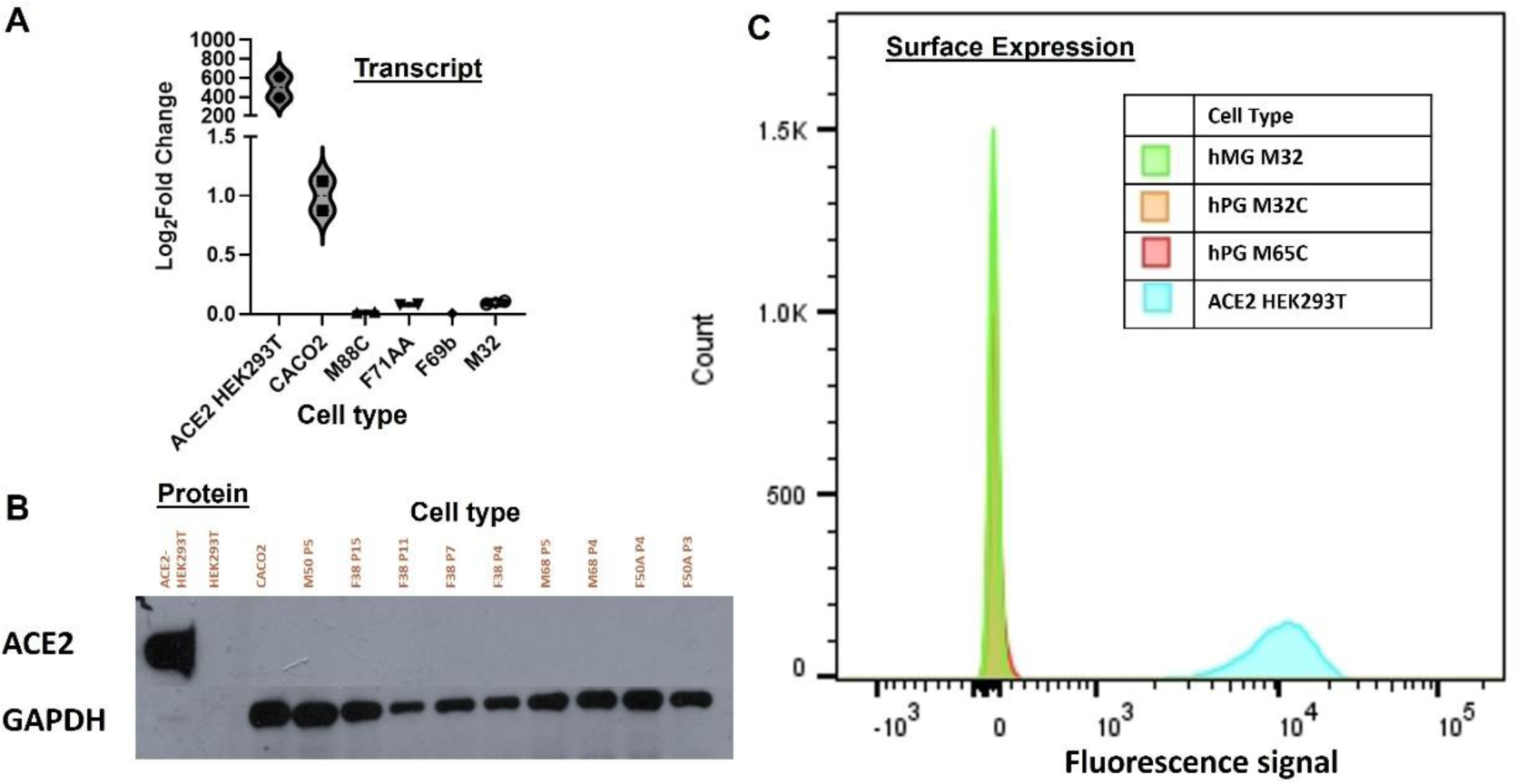
Lack of detected ACE2 transcript, protein or cell surface ACE2 expression in salivary basal cells. Three methods of detection were used to examine ACE2 levels in isolated, cultured hS/PCs equivalent to salivary basal cells. (A) qPCR results showing Log_2_Fold Change normalized to CACO2 intestinal cells. Left to right are amplimers from the ACE2 HEK293T cell line, CACO2, and four different patient-derived salivary basal cell cultures. (B) Western blot results from protein extracts from four patient-derived basal cell cultures from major and minor glands and grown to various passages. Left lane is ACE2 HEK293T cell extract followed by CACO2 and then patient-derived salivary basal cell extracts ranging from early passage (3) to late passage (15). (C) Flow cytometry analysis for ACE2 present on the surfaces of living cells. ACE2 HEK293T is shown in cyan; three salivary basal cell lines, two parotid gland-derived are shown in orange and red, and one minor salivary gland-derived is shown in green.

To examine the potential presence of ACE2 at the protein level in salivary cells, a western blot was performed using lysates from ACE2 HEK293T (positive control), HEK293T cells (negative control), CACO2 cells and expanded cultures of salivary cells from four patient donors across multiple passages to determine if ACE2 may be present in earlier passages and subsequently lost in later passages (Figure 3B). Of all lines tested, only the ACE2 HEK293T cells showed a strong positive band. Even the CACO2 cells were negative, suggesting low levels of ACE2 protein in this cell line. ACE2 was undetectable in all patient-derived salivary cells, again representing patients over a wide range of ages, different sexes and various backgrounds.

To further examine the absence of ACE2 protein on cell surfaces that might be at levels lower than could be readily detected by western blotting and to avoid the possibility that these assays were not sensitive enough to detect low levels of surface ACE2, we next conducted flow cytometry for ACE2 in primary cultured salivary cells from three individual patient donors. Again, salivary cells showed a total absence of fluorescence intensity in contrast to the positive control ACE2 HEK293T cells (Figure 3C). Taken together, these results indicated that ACE2 transcripts and protein are not detectable in primary cultures of salivary epithelial cells.

### 3.3 Spike Binding Assay on Salivary Basal Cells

Given the surprising lack of ACE2 seen in salivary tissue or cells, we sought to determine if there could be an alternative receptor on salivary cells that could bind SARS-CoV-2 spike protein to allow infection. Use of spike protein, unlike an antibody, offers an agnostic means to test the ability of the virus to bind to living cell surfaces. To verify if there was an ACE2-independent mechanism by which spike protein could bind to the surfaces of salivary cells in the absence of ACE2, we conducted flow cytometry using fluorescently labeled spike proteins corresponding to an early circulating variant of interest, alpha, and a later circulating variant of interest, omicron. We tested both spike proteins on our salivary cell populations from three patients, ACE2 HEK293T (positive control) and parental HEK293T (negative control). Flow cytometry data indicated a shift in mean fluorescence intensity on binding of spike to ACE2 HEK293T cells for both the alpha spike protein variant (Figure 4A) and the omicron spike protein variant, indicating that the spike proteins bound to cells expressing ACE2, as expected (Figure 4B). The lack of fluorescent shift indicated that basal cells derived from parotid gland were unable to bind either variant along with the unmodified HEK293T cell line.

**Figure 4.**
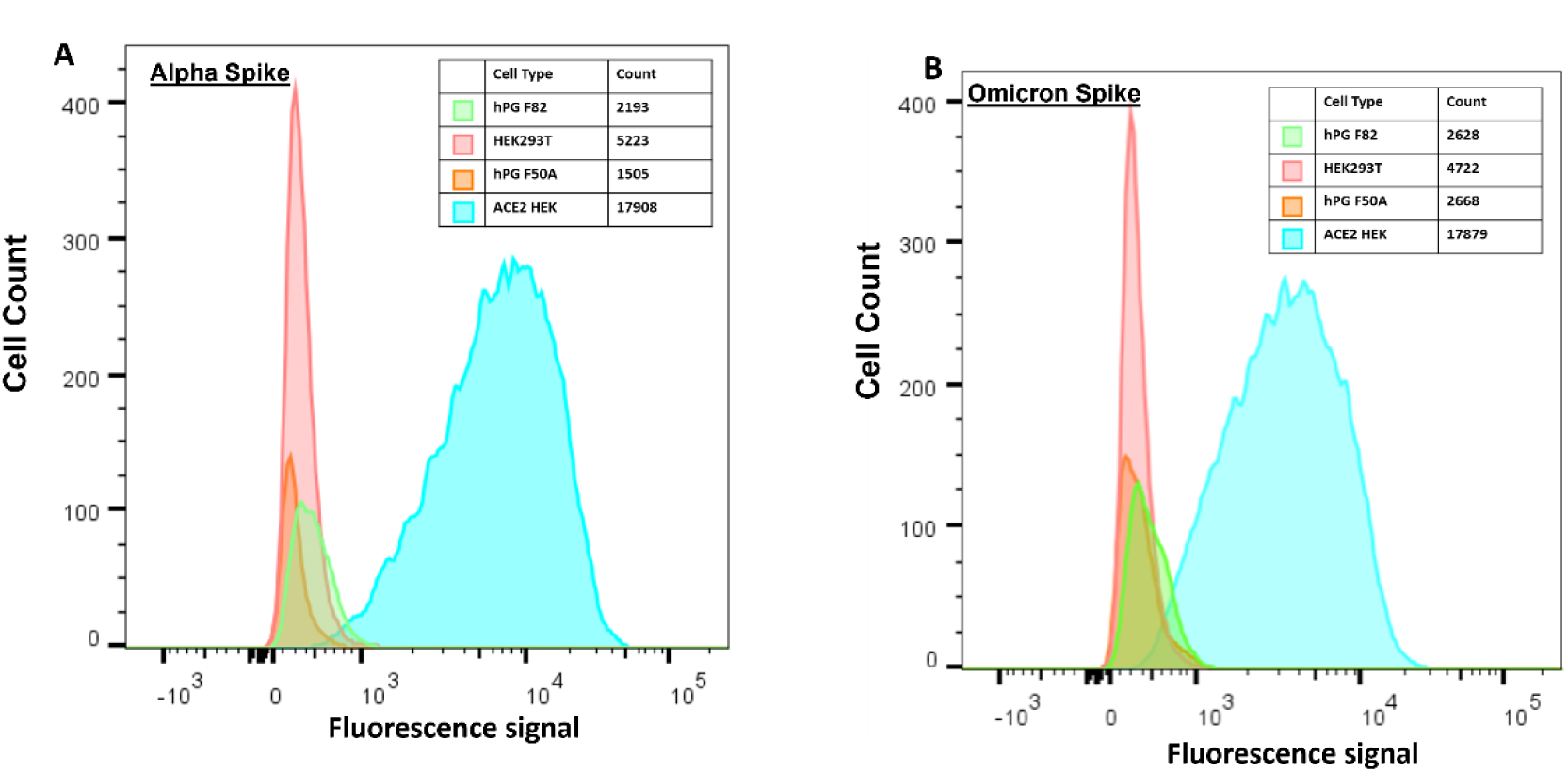
Flow cytometry to detect binding of fluorescently-labeled spike proteins corresponding to beta and omicron spike variants in living salivary cells. Two variants of spike protein were prepared as described in Methods and assessed for cell surface binding. (A) Fluorescence intensity histogram with fluorescently-labeled alpha spike protein bound to ACE2 HEK293T’s (cyan, positive control). No cell surface binding was seen for the negative control HEK293Ts (red) nor to two parotid patient-derived basal cell populations (orange and green). (B) Fluorescence intensity histogram with fluorescently-labeled omicron spike protein bound to ACE2 HEK293T’s (cyan). No surface binding was seen for HEK293Ts (red) or for two parotid patient-derived basal cell populations (orange and green).

**Figure 5.**
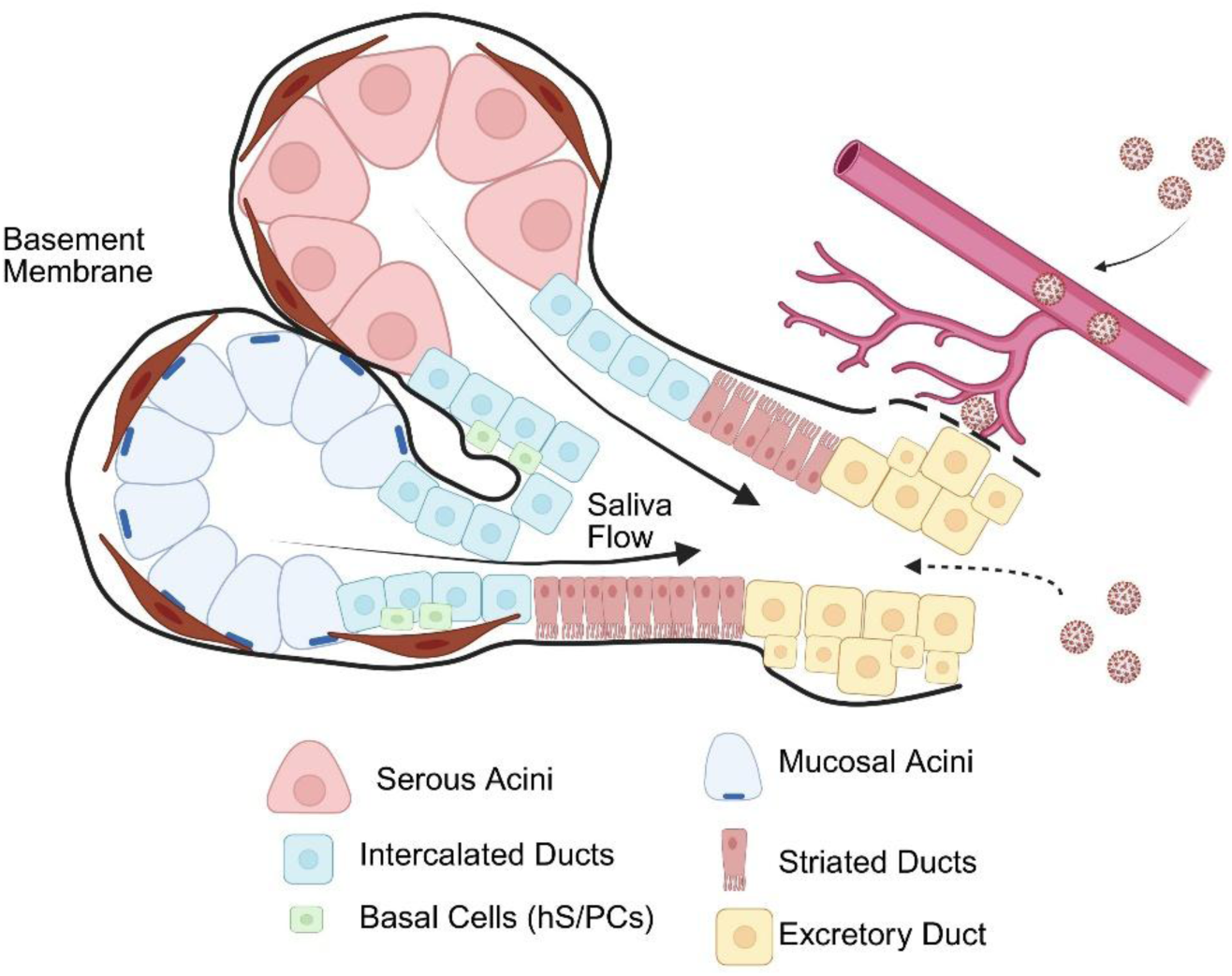
Schematic showing possible routes of major salivary gland infection by SARS-CoV-2. Saliva flow is unidirectional after secretion from the serous and mucous acini. After secretion, whole saliva is transported through the ductal network before entering the oral cavity. ACE2-dependent viral infection of major ducts through ACE2, if present on the surfaces of salivary ducts, would require retrograde ductal passage of virus from the oral cavity to reach surface receptors. The lack of detectable ACE2 suggests this is not a major route of infection. Basal cell infection would require transepithelial invasion of virus or encroachment across the basement membrane (solid and dotted black line). Salivary glands are highly vascularized; systemic SARS-CoV-2 viremia and inflammation could compromise the integrity of the basement membrane (dotted black line) allowing the virus to travel from the vasculature to the salivary gland where infection could be established in the stem/progenitor-like basal cells through an ACE2-independent route. The lack of spike binding by salivary basal cells indicates that this is not a likely alternate path of infection.

Finally, we sought to test spike binding on whole tissue sections. Formalin fixed sections and fresh-frozen sections did not show measurable GFP signal (data not shown), unsurprising since spike binding is thought to require a conformational change not possible in fixed tissue.

## 4. Discussion

Our investigation into the presence and potential involvement of ACE2 in salivary gland tissues and basal-derived salivary epithelial cells revealed a consistent lack of both transcript and protein, despite a methodologically rigorous approach that included multiple detection techniques and antibody optimization conditions. The lack of ACE2 RNA transcripts and protein levels in basal cells derived from both parotid and minor salivary glands (Fig 1) was surprising given that the literature indicates that basal ductal cells were likely to have ACE2 [27], [35]. These findings directly contradict some earlier studies and suggest that salivary glands are unlikely to serve as major entry points or reservoirs for SARS-CoV-2 via an ACE2-dependent route.

Our first studies evaluated ACE2 protein expression at the tissue level in both parotid and minor salivary glands using immunocytochemistry. To address variability seen in the literature, we systematically tested three well-characterized ACE2 antibodies under a range of antigen retrieval conditions (no retrieval, pH 6, and pH 9). Our positive control, small intestinal tissue, exhibited robust apical ACE2 staining under pH 6 and pH 9 retrieval conditions, confirming the efficacy of the antibodies and the staining protocol. In contrast, salivary tissues showed no ACE2 signal under any condition, despite the clear labeling of epithelial (pan-cytokeratin) and myoepithelial (α-SMA) cells. These results indicate that neither the acinar nor ductal compartments of the parotid and minor salivary glands express detectable ACE2 protein.

To validate these findings with a complementary approach, we performed western blotting on lysates from both parotid and minor salivary glands. Consistent with the immunostaining data, ACE2 protein was undetectable in all salivary gland samples, whereas it was readily observed in small intestine lysates and ACE2-overexpressing HEK293T cells. These results support the conclusion that ACE2 is not expressed at significant levels in exocrine salivary cells under basal conditions.

Given the potential for disease-induced upregulation or the presence of receptor expression below detection thresholds in tissue sections, we extended our investigation to primary human salivary epithelial cell cultures (hS/PCs) derived from both major and minor glands of multiple patients across sex, age, and ethnic backgrounds. Quantitative RT-PCR showed that none of these cell lines expressed detectable ACE2 transcripts, even after treatment with the pro-inflammatory cytokines TNFα and IFNγ for 24 and 72 hours.

Furthermore, western blot analysis of four different donor-derived cell cultures across multiple passages, again showed no ACE2 protein, and this was confirmed with highly sensitive flow cytometry. Across all patient-derived lines, there was no detectable ACE2 surface expression, while ACE2-overexpressing HEK293T cells consistently served as a strong positive control.

While ACE2 is the primary receptor for SARS-CoV-2, there have been several publications that have implicated additional receptors for the virus to bind and enter through integrins [29], [36]. To explore the possibility of ACE2-independent viral binding mechanisms, we used spike proteins corresponding to two SARS-CoV-2 variants (Alpha and Omicron) labeled for fluorescence to probe basal salivary cells via flow cytometry. This method provides a direct, receptor-agnostic approach to assess potential viral engagement with the cell surface. However, no binding was observed in salivary cells from any patient donor, in contrast to the strong fluorescent shifts seen in ACE2-expressing HEK293T cells. These data indicate that neither canonical ACE2-dependent nor alternative receptor-mediated spike binding occurs in basal salivary epithelial cells under the conditions tested.

These findings have significant implications for understanding viral interactions in the oral cavity. Early in the COVID-19 pandemic, public transcriptomic databases and single-cell RNA sequencing data suggested the salivary glands might express ACE2, leading to speculation that ducts could serve as viral portals or reservoirs [9], [27], [37]. However, those datasets showed only low, cell-type-specific ACE2 mRNA expression, and many lacked robust validation at the protein level. Our data indicate that even under optimized conditions, salivary epithelial cells lack both transcript and protein expression of ACE2. While some papers have shown ACE2 protein in the ducts [38] or both acinar cells and ducts [39], [40], the literature routinely shows inconsistency in immunocytochemistry results as well as a robust protein expression that does not correlate with the transcript levels. Additional studies supported our results suggesting that the salivary glands do not have ACE2 at the protein level [7] and another study suggested that the inconsistency between these data may be a result of the overlap in homology between ACE2 collectrin-like binding domain [41]. The inconsistency across the literature may partly stem from the use of antibodies with cross-reactivity to domains with high homology to the ACE2 such as collectrin, potentially leading to false-positive immunostaining results.

Moreover, our findings argue against models proposing retrograde infection of the salivary ducts from the oral cavity as a significant route of SARS-CoV-2 entry. If such a mechanism were valid, one would expect to observe ACE2 on ductal surfaces or successful spike protein binding in vitro, neither of which was observed in our study.

While our results refute local epithelial susceptibility to SARS-CoV-2 via ACE2 in the salivary glands, we acknowledge that glandular involvement during systemic viral infection remains plausible through hematogenous spread. Salivary glands are highly vascularized, and several viruses, such as mumps and CMV [42], can infect glandular tissues via the bloodstream [43], [44]. In such cases, the infection may involve infected perivascular, immune or stromal cells rather than epithelial compartments. SARS-CoV-2 may similarly affect salivary tissues through systemic routes, potentially explaining the detection of viral RNA in salivary secretions in some patients [27], albeit without clear evidence of productive infection in glandular epithelial cells.

In summary, our systematic study clearly demonstrates that ACE2 is absent in basal cells and whole tissue from both major and minor salivary glands, and that these cells are not likely to be susceptible to initial infection of SARS-CoV-2 via spike binding, even through alternative pathways. These results collectively challenge prior assumptions regarding the role of salivary glands in direct SARS-CoV-2 pathogenesis and highlight the importance of multimodal validation in receptor expression studies of infection.

## 5. Conclusions

In conclusion, our data, together with the anatomical context of major and minor salivary gland architecture and position, support the hypothesis that direct infection of salivary glands via the oral cavity is a highly improbable first route of infection. The physical barriers posed by the mucosal layer, the protective properties of saliva, and the complexity of the ductal system, including Stensen’s and Wharton’s ducts, make retrograde viral entry and surface infection via ACE2 an unlikely route. Instead, we propose that in some cases, SARS-CoV-2 reaches the salivary glands through systemic dissemination, gaining access via the glandular vasculature, immune cells or stroma. This idea is consistent with the expression patterns and levels of ACE2 in cells and tissues and may help explain observed viral presence in glandular tissues including salivary glands of infected patients. Furthermore, the potential for spike-independent mechanisms of entry cannot be excluded, especially in cellular contexts where ACE2 is minimally expressed or absent. Such alternate pathways, possibly involving endosomal uptake or alternative receptors, may expand the tissue distribution of SARS-CoV-2 and warrant further investigation in glandular models. These insights contribute to a broader understanding of viral pathogenesis within the oral cavity and underscore the importance of considering local cellular susceptibility in determining infectivity.

## Abbreviations

The following abbreviations are used in this manuscript:

SARS-CoV-2: Severe Acute Respiratory Syndrome Coronavirus 2

ACE2: Angiotensin Converting Enzyme 2

hS/PCs: human stem/progenitor cells

α -SMA: α-smooth muscle actin

Pan CK: pan-cytokeratin

## Author Contributions

Conceptualization, C.B., S.Y. and MC.F-C.; methodology C.B.; formal analysis C.B., S.Y. and M.C.F-C.; resources S.Y. and M.C.F-C.; data curation C.B.; writing-original draft preparation C.B. and M.C.F-C.; writing-review and edition C.B.; S.Y.; M.C.F-C.; supervision S.Y. and M.F-C.; project administration S.Y. and M.F-C.; funding acquisition S.Y. and M.F-C. All authors have read and agreed to the published version of the manuscript.

## Institutional Review Board Statement

The study was conducted in accordance with the Declaration of Helsinki, and approved by the Institutional Review Board of University of Texas Health Science Center of Houston (HSC-DB-17-0855 approved 10-30-2017) and (HSC-DB-20-0720 approved 07/15/2020). No animals were involved in this study.

## Informed Consent Statement

Informed consent was obtained from all surgical subjects donating tissues used in the study. All patient data was deidentified for this study except for age and sex and no patients can be identified from the information provided in the study.

## Data Availability Statement

Data available upon reasonable request.

## Acknowledgments

We would like to acknowledge Dr. Roy Davee and Dr. Jaiyeola Thomas-Ogunniyi for the initial small intestine tissue that allowed for the validation of our antibody panel. Dr. Robert Witt (Christiana Care, Newark DE) and Drs. Andrew Farach and Nadia Nadia G. Mohyuddin (Houston Methodist Hospital) for major salivary gland tissues. We would also like to acknowledge Dr. Daniel Carson for his input on the data during lab meetings that helped spark the conversation to identify ACE2-independent methods for spike binding as well as the moral support to CB provided in tackling experiments that seemingly consistently contradicted the literature. Confocal fluorescence microscopy images and the BCA assays on cytation 5 plate reader were obtained at Center for Craniofacial Research Instrumentation Core, School of Dentistry, UTHealth Houston. The flow cytometry experiments were run at the Flow Cytometry and Cellular Imaging Core Facility, the University of Texas MD Anderson Cancer Center, Houston. Sample preparation including paraffin embedding and sectioning were done by the Oral and Maxillofacial Pathology Core at UTHealth Houston School of Dentistry. The authors thank all the members of the Farach-Carson and Young labs for their many helpful discussions during the course of these studies and in particular, Drs. Danielle Wu and Neeraja Dharmaraj for their generosity in sharing their expertise in all endeavors.

