## Supplementary Figure 1 and Table 1 for "Exploring Spike-Dependent and ACE2-Independent Viral Entry into Salivary Epithelial Cells in the Absence of ACE2"

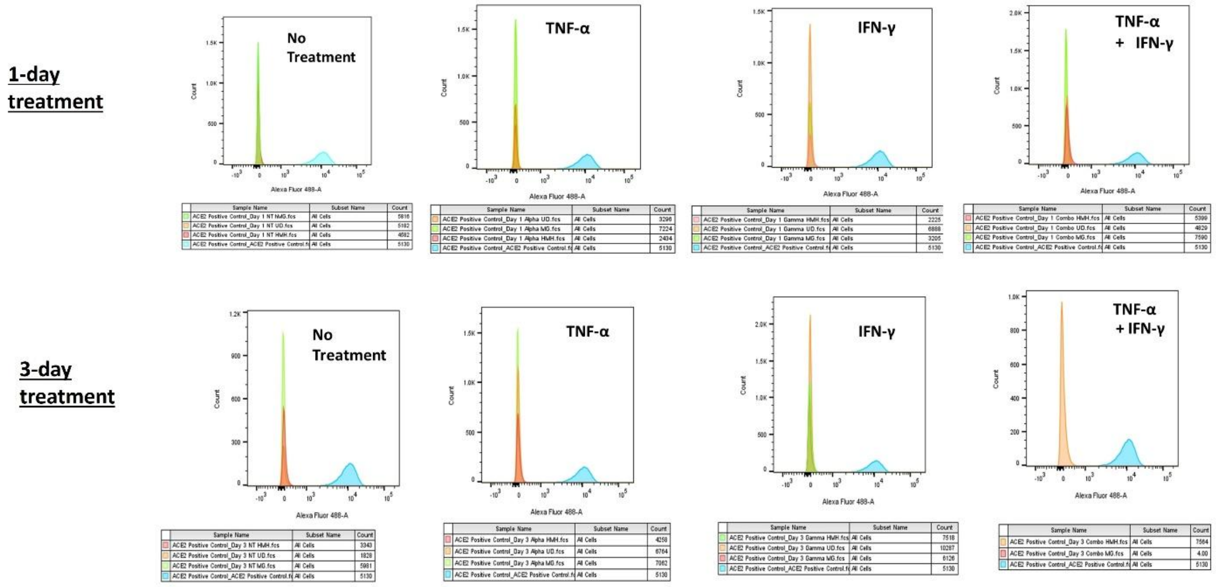

**Supplementary Figure 1. Cytokine treatment fails to induce ACE2 protein expression in salivary cells as assessed by flow cytometry**. Cells were treated with inflammatory cytokines for various times, harvested and analyzed for ACE2 cell surface expression by flow cytometry as described in the Methods. (A) 24 hours of treatment. (B) 72 hours of treatment. Left to right: No treatment, TNF-α, IFN-γ and combination of the two.

### Supplemental Table 1. Antibodies Used in this Study

| Antibody | Target | Description | Source |
|---|---|---|---|
| MAb 66699 | ACE2 | mouse monoclonal | Protein Tech |
| MAb 272500 | ACE2 | mouse monoclonal | Abcam |
| PAb 15348 | ACE2 | rabbit polyclonal | Abcam |
| MAb7817 | SMA | mouse monoclonal | Abcam |
| MAb234297 | Pan-cytokeratin | rabbit monoclonal | Abcam |
| A11008 | anti-rabbit | goat polyclonal (488) | Invitrogen |
| A11004 | anti-mouse | goat polyclonal (568) | Invitrogen |
| A11029 | anti-mouse | goat polyclonal (488) | Invitrogen |
| A11036 | anti-rabbit | goat polyclonal (568) | Invitrogen |
